# Combined inhibition of NAD synthesis and C-terminal binding protein cooperatively induce cell death and inhibit growth of High Grade Serous Ovarian Carcinoma

**DOI:** 10.1101/2025.03.22.643181

**Authors:** Kranthi Kumar Chougoni, Jacqueline West, Nicholette St. Martin, Diana Dcona, Istem Kose, Nari Kim, Martin M. Dcona, Ronny Drapkin, Joseph W. Carlson, Ann E. Walts, Sandra Orsulic, Keith C. Ellis, Steven R. Grossman

## Abstract

The transcriptional scaffolds C-terminal Binding Proteins (CtBP) 1 and 2 are overexpressed and act as oncogenic dependencies in multiple cancers but importantly encode a chemically targetable dehydrogenase domain. CtBP promotes survival of high grade serous ovarian carcinoma (HGSOC) cells by repressing expression of Death Receptors (DR) 4 and 5, which activate caspase 8-dependent apoptosis. We have previously developed a series of substrate competitive CtBP dehydrogenase inhibitors active in multiple cell and preclinical solid tumor models. In the current study, we validated CtBP 1 and 2 overexpression in a longitudinal series of primary and metastatic/recurrent HGSOC cases. Furthermore, our lead CtBP dehydrogenase inhibitor, JW-98 induced apoptosis and exhibited variable single agent IC_50_ values in HGSOC cell lines, but depletion of nicotinamide adenine dinucleotide (NAD) using the NAD synthesis inhibitor GMX1778 strikingly sensitized tumor cells to JW-98 treatment. Mechanistically, the JW-98/GMX-1778 combination effectively disrupted CtBP dimerization that requires stoichiometric levels of intracellular NAD and is required for oncogenic transcriptional activities. Highlighting the translational potential of this combination, combined JW-98/GMX1778 treatment of OVCAR3 HGSOC xenografts in immunodeficient mice abrogated tumor growth without observable toxicity. CtBP/NAD combined inhibition represents a novel therapeutic strategy that could improve outcomes in chemoresistant HGSOC.

## Introduction

Epithelial Ovarian Cancer (EOC) afflicts nearly 20,000 women each year and accounts for more than 12,000 cancer deaths among US women annually (1). The most common EOC subtype, High Grade Serous Ovarian Cancer (HGSOC), is an especially lethal cancer due to a lack of early detection strategies resulting in diagnosis at an advanced stage, frequent development of chemoresistance to standard of care therapies, and lack of targetable driver oncogenes (2). Thus, novel therapeutic strategies for advanced/refractory disease have focused on targetable vulnerabilities and dependencies within oncogenic pathways that distinguish HGSOC cells from normal cells. We identified the C-terminal binding protein (CtBP) family of transcription factors, of which CtBP2 was already known to be overexpressed in the majority of EOC tumors (3), as oncogenic drivers that promote survival of HGSOC cells by repressing expression of the proapoptotic Death Receptors 4/5 (4). Functionally, CtBP1 and 2 are paralogous dehydrogenases and transcriptional scaffolds/coregulators, and their overexpression in many solid tumors uniformly correlates with worse prognosis (5). CtBP and its interactome contribute to malignant progression by repressing proapoptotic (DR4/5, Bik, Bax) (5), tumor-suppressor (E-cadherin, PTEN) (5), and activating oncogenes (Tiam1, c-Myc) (5, 6). Furthermore, in both pancreatic and colon cancer mouse models, CtBP drives tumor progression and metastasis by promoting cancer stem cell activity (6, 7).

We have developed a library of CtBP dehydrogenase substrate competitive inhibitors (CtBPi) centered around the scaffold of the 1st generation CtBP inhibitor hydroxyimino-3-phenylpropanoic acid (HIPP) that are active across a spectrum of solid tumor types, including colon and pancreatic cell lines and mouse models. Our 2nd generation CtBPi, 4-chloro-HIPP (4-Cl-HIPP), phenocopied the hypomorphic effects of *Ctbp2* allelic deletion in *Apc min* mice, attenuating intestinal polyposis and extending survival (7). As nicotinamide adenine dinucleotide hydride (NADH) is embedded in CtBP’s ternary structure, we have further hypothesized that limiting the cellular concentration of NAD species (NAD+/NADH) might enhance efficacy of CtBP inhibitors. Indeed, GMX1778, an inhibitor of the NAD synthetic enzyme nicotinamide phosphoribosyl transferase (NAMPT), synergized with 4-Cl-HIPP to kill pancreatic cancer cells in culture and strongly limit growth of pancreatic cancer xenograft tumors (8).

In this current work we demonstrate the cytotoxic efficacy in HGSOC cells and tumors of a novel 3^rd^ generation CtBP inhibitor JW-98, in which the carboxyl moiety of 4-Cl-HIPP has been esterified to improve cell penetration. Furthermore, limiting cellular NAD synthesis with GMX1778 significantly enhanced sensitivity of the tumor cells to cell death induced by CtBP inhibitor (8).

Upon mechanistic investigation, we found stoichiometric disruption of CtBP dimers upon combined treatment with GMX1778 and JW-98, providing a proof-of concept for utilizing CtBP/NAMPT inhibitor combinations in HGSOC. When tested for therapeutic utility using an HGSOC xenograft mouse model, CtBP/NAMPT inhibitor combination treatment abrogated tumor growth, demonstrating the potential clinical utility of this combination. Overall, our new findings suggest combining NAD depleting agents with CtBP inhibitors could be an effective strategy to safely and effectively target HGSOC.

## Materials and Methods

### Cell lines and Reagents

OVCAR 3, 4, 8, OVCA429, and Kuramochi HGSOC cells (4, 9), were grown in 10 % DMEM with 1% penicillin and streptomycin. All cell lines were routinely tested for mycoplasma (Abcam, Cat No: ab289834).

### JW-98 synthesis

A Horner-Wadsworth-Emmons (HWE) reagent was synthesized (**Fig S1, Scheme 1**) by first coupling dimethyl phosphite (1 eq) and ethyl glyoxylate (1 eq) with triethylamine (TEA, 0.2 eq) in dichloromethane (DCM) at -78°C for 1 hour to afford ethyl 2-(dimethoxyphosphoryl)-2-hydroxyacetate (**Fig. S1, 1.1**). This was then TBS protected with tert-butyldimethylsilyl chloride (2 eq), imidazole (3 eq), and DMAP (0.15 eq) in DCM at room temperature overnight to yield the final HWE reagent, ethyl 2-((tert-butyldimethylsilyl)oxy)-2(dimethoxyphosphoryl)acetate (**Fig. S1, 1.2**). 4-chlorobenzaldehyde scaffold was then subjected to a HWE reaction in the presence of LiHMDS (1.1 eq), where the aldehyde was used in excess (1.1 eq) relative to the HWE reagent (1 eq). The HWE reaction was carried out with LiHMDS as a base under reflux overnight in tetrahydrofuran (THF) following (**Fig. S1, Scheme 2)**. After purification, the resulting silyl enol ether (**2.1**) underwent a two-step, one-pot reaction in which the silyl enol ether is TBS de-protected using triethylamine trihydroflouride (1.7 eq), and the resulting enol is converted to an oxime with hydroxylamine hydrochloride (1.7 eq) to yield the final oxime product in room temperature conditions overnight in two parts chloroform, and one part ethanol (**Fig. S1, 2.2**). Average purity was assessed by LC-MS was 95%.

### Cell Viability Assays

3 × 10^3^ OVCAR 3, 4, 8, OVCA429, or Kuramochi cells were seeded in 96 well plates, and after overnight incubation, treated with vehicle or JW-98 for 72 hours, after which cell viability was determined by crystal violet staining. Absorbance was measured at 590 nm after dissolving the crystal violet stain in 10% glacial acetic acid and IC_50_ values were calculated using GraphPad Prism.

### Drug Combination Assays

Equal number of HGSOC cells were seeded in 6 well plates. After an overnight incubation, cells were pretreated with NAMPT inhibitor GMX1778 for 24 h to deplete NAD levels following the treatment with JW-98. Cell viability was assessed by crystal violet staining as described above.

### Cross-linking assay

Disuccinimidyl glutarate (DSG) cross-linker (Thermo Fisher Scientific, #20593) was prepared as a stock solution of 100 mmol/L in DMSO made fresh for each experiment. *In vivo* cross-linking was carried out in OVCAR 3 cells that were collected by scraping, washed with cold PBS (pH 7.2), and resuspended in PBS (pH 8.2) with 1X Complete Protease Inhibitor Mixture, EDTA-free (Roche). Samples were incubated with cross-linker (1 mmol/L) for 30 minutes at 37°C with rotation. The reaction was quenched with the addition of 1 mol/L Tris HCl pH 7.6 to 20 mmol/L final concentration and incubated for 15 minutes at room temperature. After quenching, *in vivo* cross-linked samples were lysed using RIPA Lysis and Extraction Buffer (Thermo Fisher Scientific, #89900) followed by ultracentrifugation for 30 minutes at 15,000 rpm at 4°C. Samples were then immunoblotted for CtBP2.

### Annexin V and Propidium Iodide staining

2 × 10^6^ OVCAR3 cells were seeded in a 15 cm dish, after an overnight incubation cells were treated with indicated JW-98 drug concentrations (0, 50, 100, and 200 µM). 5 days following drug treatment, attached and floating cells were collected and counted using trypan blue exclusion assay. 1 × 10^6^ cells were stained using Alexa Fluor 488 Annexin V/Dead Cell Apoptosis Kit as per manufacturer instructions (Thermofisher Scientific, Cat No: V13242). Flow cytometry was performed using BD X Fortessa 20, and percentage of apoptotic cells (Annexin V/PI positive cells) was quantified after recording 20000 events.

### NAD/NADH measurement

3 × 10^3^ OVCAR3 or OVCA429 cells were seeded in a 96 well plate. After overnight attachment of cells, the cells were treated with GMX1778 (0 or 1 nM) for 48h following which the NAD levels were assessed using a Promega-glo NAD/NADH measurement kit (Cat No : G9071) as per manufacturer instructions.

### Xenograft study

2-month-old NSG female mice were subcutaneously injected with 2 × 10^6^ OVCAR3 cells. 10 days after tumor cell injection, mice were randomly assigned to one of the treatment groups with either vehicle, JW-98 (100 mg/Kg), GMX1778 (30 mg/Kg) or JW-98 and GMX1778 for two weeks. Tumor volumes were measured weekly, and tumor weights were calculated at the end of the study. All the animal studies were approved by USC Institutional Animal Care and Use and Committee.

### Immunohistochemistry

Formalin-fixed paraffin-embedded (FFPE) tissue sections were deparaffinized and dehydrated, following which antigen unmasking was performed using Retriever 2100 (Aptum Biologics Ltd). Slides were blocked in 5% Goat serum (Thermo Fisher Scientific, Cat No : 31872) for 1 h at room temperature and incubated with primary antibody (CtBP1 (Cat No: 612042, mouse anti-CtBP1; 1:25; BD Biosciences), CtBP2 (Cat No: 612044, mouse anti-CtBP2; 1:25; BD Biosciences), Cleaved Caspase 3 (Cat No: 9661; rabbit anti-cleaved caspase 3; 1:100; Cell Signaling Technology)) overnight at 4º C. Following three 5 min washes in 1X PBST, the slides were then incubated in secondary antibody (Goat Anti-Mouse IgG (H + L)-HRP Conjugate mouse, dilution, 1:200, Cat No: 1706516; Goat Anti-Rabbit IgG (H + L)-HRP Conjugate rabbit, dilution, 1:200, Cat No :1706515; Bio-Rad Laboratories) for 1 h at room temperature and stained using DAB substrate (DAKO Chromogen, Agilent) per manufacturer instructions. Nuclei were then counterstained using Mayer’s Hematoxylin (Cat No: 2617303, Electron Microscopy Sciences) for 3 min, and slides were dehydrated and cover slipped as previously described (10). The intensity of CtBP1/2 nuclear staining was scored on a scale 0-3; 0 for no staining, 1+ for weak staining, 2+ for moderate staining and 3+ for strong staining.

### Statistical Analysis

All the results presented represent an average of at least three independent experiments. All the pairwise comparisons between two groups were performed using paired student t test using GraphPad Prism software.

## Results

### Expression of CtBP1/2 across the HGSOC disease spectrum

Multiple studies have shown overexpression of both CtBP1/2 proteins in various solid tumors to be linked to poor outcomes (5). A prior study analyzed CtBP2 expression in an EOC series that included a limited number of HGSOC cases, but CtBP2 expression specifically in HGSOC was not reported (3). We therefore analyzed a patient-matched longitudinal set of HGSOC specimens for CtBP1/2 expression by immunohistochemistry (IHC; **Fig. 1**). The series of 42 cases includes primary tumors and concurrent metastases obtained at primary debulking surgery, as well as recurrent tumors obtained at second-look surgeries after treatment (usually platinum/taxane combination) (11). Notably, CtBP1 was uniformly highly expressed at all stages (**Fig. 1B, 1D**), while CtBP2 staining was more varied in primary tumors, but demonstrated more universal expression in advanced stages (**Fig. 1C, 1E**). By contrast, normal fallopian tube (FT) samples were negative or borderline detectable for CtBP2 (**Fig. 1A**). Thus, HGSOC tumors might initially select for elevated expression of CtBP1, and with malignant progression could further select for elevated expression of CtBP2.

**Fig. 1.**
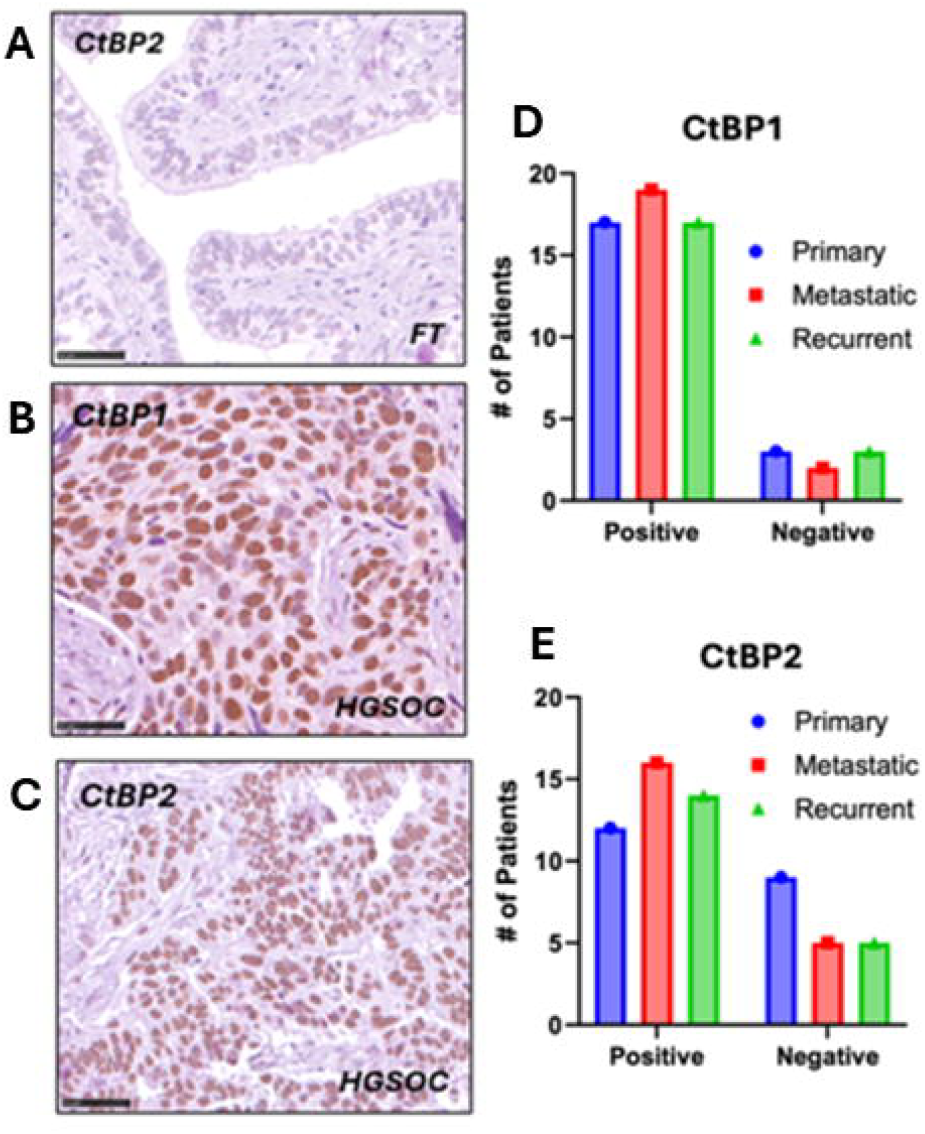
Expression of CtBP in HGSOC. **(A-C)** IHC using CtBP1/2 antibodies performed on sections of resected normal fallopian tube (FT; **A**) and primary HGSOC (**B-C**). **(D-E)** Quantitation of CtBP1/2 IHC staining from a series of HGSOC cases representing intra-patient primary, metastatic, and recurrent tumor. Positive indicates 1+ to 3+ staining. Error bars indicate +/- 1.0 SD. Scale bar = 50 µM.

### Development of a novel 3^rd^ generation CtBP inhibitor

We have iteratively improved the potency of CtBP inhibitors (12). Our 1st generation inhibitor, hydroxyimino-3-phenylpropanoic acid (HIPP), was rationally designed from substrate/catalytic domain interactions observed in the co-crystal structure of CtBP with its substrate 2-keto-4-methylthio-2-oxo butyric acid (MTOB) and based on the dehydrogenase enzymatic mechanism (12). We then identified the potent 2^nd^ generation CtBP inhibitor 4-Cl-HIPP based on a structure-activity relationship study of substituents on the HIPP phenyl ring (7). In developing the 3rd generation of CtBP inhibitors, we sought to improve cell-permeability through esterification of the 2nd generation CtBPi 4-Cl-HIPP to form our lead CtBPi JW-98 **(Fig. 2)**.

**Fig 2.**
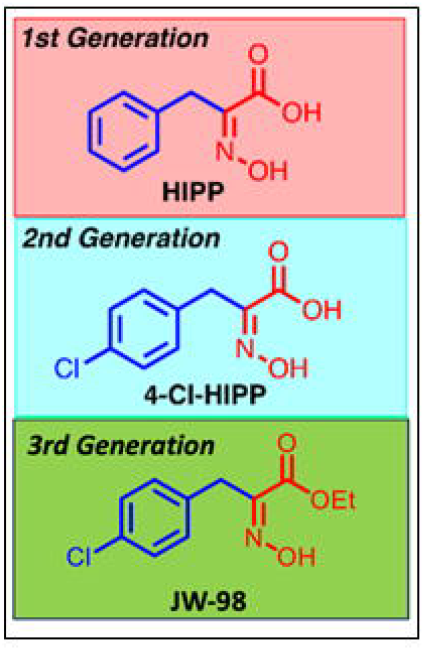
Chemical structures of 1^st^-3^rd^ generation HIPP-class CtBP substrate-competitive dehydrogenase inhibitors. JW-98 is the ethyl ester of 4-Cl-HIPP.

### JW-98 induces cell death by apoptosis in HGSOC cells

Our previous work in HGSOC cells demonstrated repression of pro-apoptotic genes DR4/5 by CtBP in HGSOC cells (4). Furthermore, transient or stable knockdown of CtBP1/2 led to cell death by Caspase 8 dependent apoptosis, unveiling the CtBP dependency of HGSOC cells. To determine if a CtBP inhibitor could phenocopy genetic CtBP loss in HGSOC, we assessed the cytotoxic efficacy of JW-98 in a panel of HGSOC cell lines (OVCAR3, OVCAR4, OVCAR8, OVCA429 and Kuramochi) including platinum resistant (OVCAR3, OVCAR8) cell lines (13). Upon titrating increasing JW-98 concentrations into HGSOC cell lines over a period of 7 days, we observed measurable IC_50_’s in the vicinity of ∼100 µM in some cell lines (OVCAR3, OVCAR4, Kuramochi), while OVCAR8 and OVCA429 cells were resistant to JW-98 **(Fig. 3A)**. Thus, despite esterification of 4-Cl-HIPP, the IC_50_ values of JW-98 in HGSOC cells as a single agent were higher than optimal and suggested that further development might require combination strategies to enhance efficacy at lower doses.

**Fig 3.**
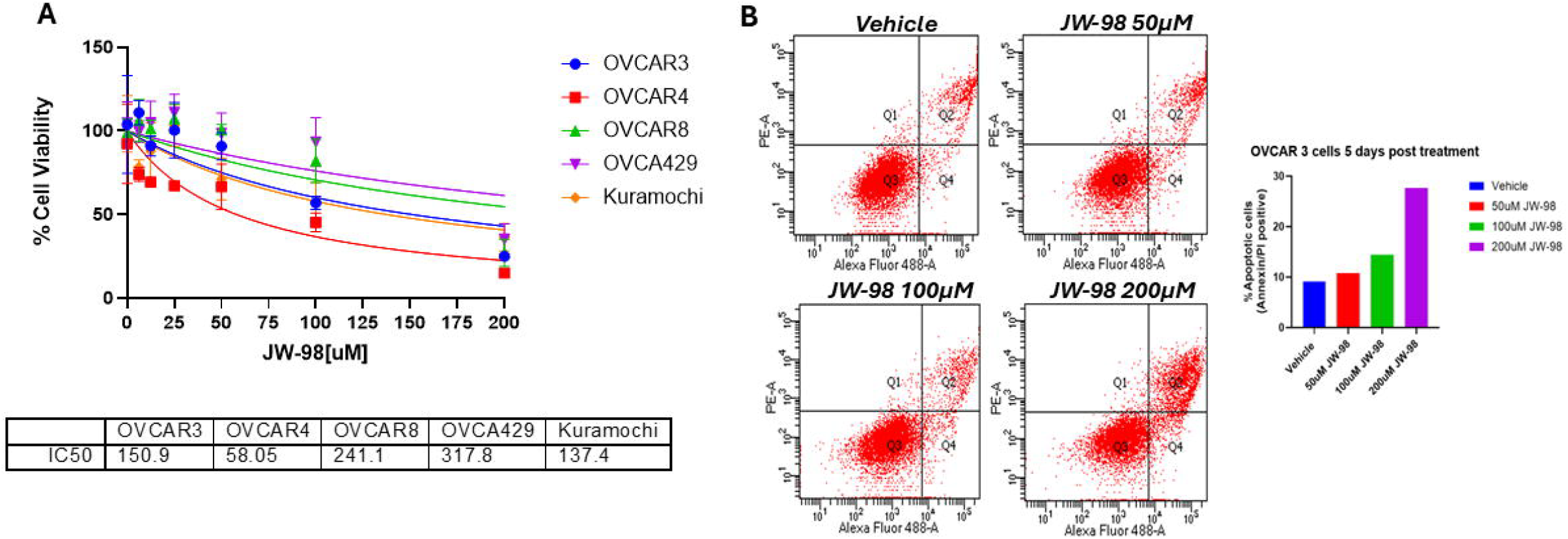
Cytotoxicity induced by JW-98 in HGSOC cells. **A**. The indicated cell lines were treated with vehicle or increasing doses of JW-98 for 72 h, cell viability was assessed by crystal violet staining, with absorbance of solubilized dye recorded at 590 nm. IC_50_ values were calculated using GraphPad Prism (N=3). Error bars indicate +/1.0 SD. **B**. OVCAR3 cells were treated with vehicle or JW-98 for 5 days and analyzed by Annexin V-488/Propidium Iodide staining. The percentage of apoptotic cells was quantified using flow cytometry.

### Mechanism of cell death induced by JW-98

Next, we probed for mechanism of cell death induced by JW-98 in cells with single agent sensitivity. It has been demonstrated in HGSOC cells that genomic depletion of both CtBP1 and CtBP2 leads to apoptosis (4). Hence, we analyzed JW-98 treated cells with Annexin V and propidium Iodide staining followed by flow cytometry to assess the induction of apoptosis. Notably, JW-98 treatment showed a significant increase in apoptotic cells (∼30%) in a dose dependent manner compared to vehicle treatment **(Fig. 3B)**. Thus, CtBP chemical inhibition induces HGSOC cell apoptosis as does depletion via RNAi (4).

### JW-98 along with GMX1778 disrupts oligomeric CtBP complex

We and others have shown CtBP activity is dependent on the availability of NAD for oligomerization or dimerization to form transcriptional complexes (14). NAD in cancer cells is synthesized using either the salvage or Preiss-Handler pathways, which are dependent on the NAMPT or NAPRT enzymes, respectively, though only NAMPT chemical inhibitors are currently commercially available. Moreover, CtBP oligomerization status in a cell can serve as a biomarker for activated oncogenic transcriptional programs (15). We have previously demonstrated disruption of cellular CtBP dimerization by 4-Cl-HIPP in pancreatic cancer cells in the setting of NAD depletion induced by the NAD synthesis inhibitor GMX1778, which is a potent inhibitor of NAMPT (8).

To establish if JW-98 similarly disrupts CtBP dimerization in HGSOC, we first established whether GMX1778 treatment could deplete NAD levels in OVCAR3 cells. Indeed, treatment of OVCAR3 cells with a non-toxic dose of GMX1778 strongly depleted intracellular NAD levels (NADH/NAD+ species), which was rescued by addition of NAD synthesis precursor nicotinic acid demonstrating that NAD depletion by GMX1778 was due to on-target activity **(Fig 4A)**. We next assayed CtBP dimerization in OVCAR3 cells treated with JW-98, GMX1778, or the combination, by treating live cells with disuccinimidyl glutarate (DSG) crosslinker followed by immunoblotting of lysates for CtBP2 **(Fig. 4B)**. Strikingly, neither drug alone reduced dimerization, but the combination potently reduced and even eliminated dimerization in a dose dependent manner for JW-98 (**Fig. 4C**).

**Fig 4:**
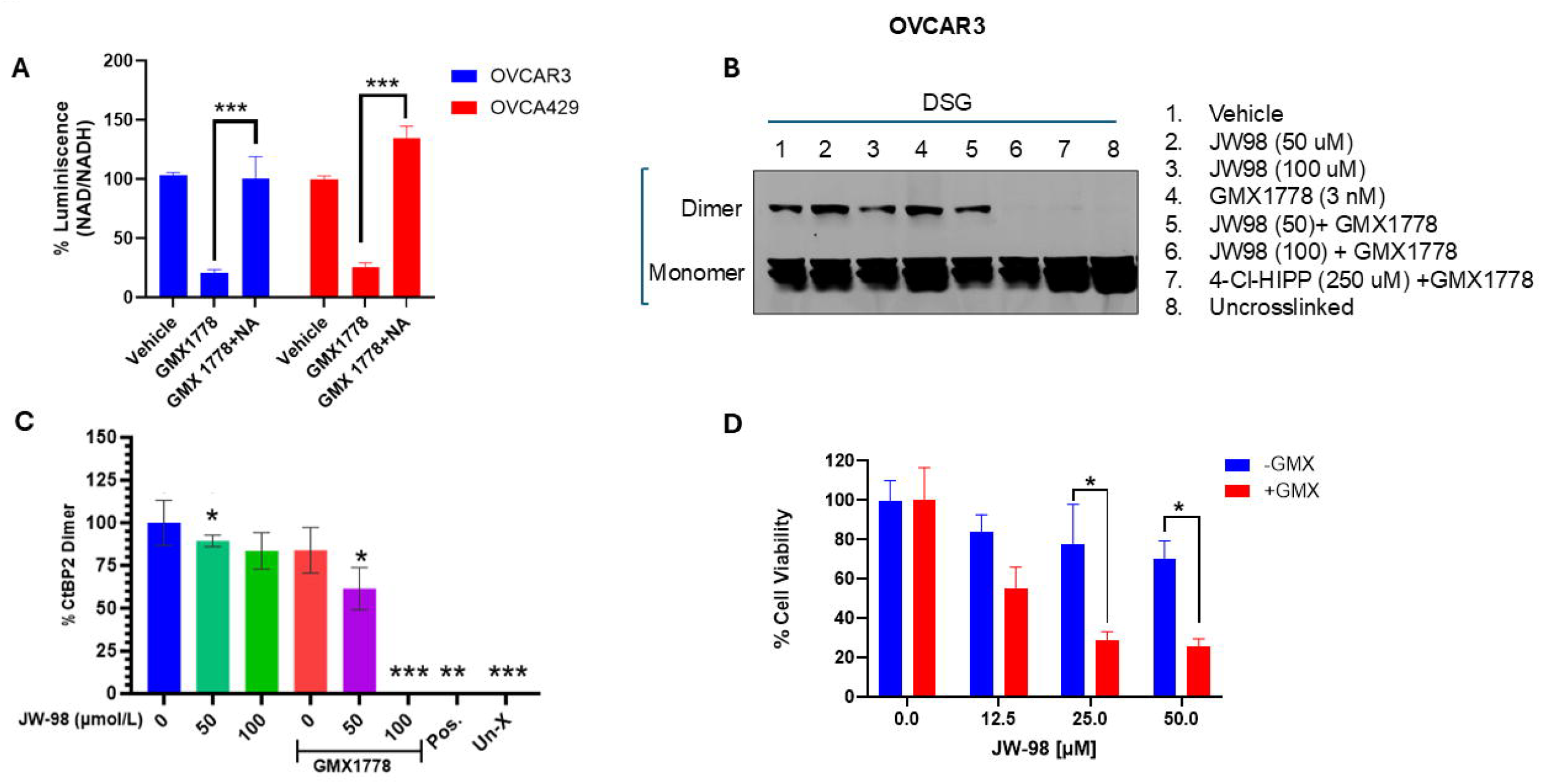
NAMPT inhibitor/JW-98 combination inhibits CtBP oligomerization and induces cytotoxicity in HGSOC cell lines. **A**. NAD/NADH levels in OVCAR3 were measured 48 hours after GMX1778 treatment using NAD/NADH assay kit (Promega-glo) and % NAD depletion was quantified by measurement of luminescence. Nicotinic acid NAD rescue was used as a control for on-target specificity. **B**. OVCAR3 cells were treated with Vehicle (lanes 1-3) or 3 nmol/L GMX1778 (lanes 4–6) for 24 hours, followed by addition of Vehicle (0) or indicated concentrations of 4-Cl-HIPP for 48 hours, followed by incubating live cells in 0.25 µmol/L DSG. Lane 7 represents a positive control using 4-Cl-HIPP/GMX1778 combination. Lane 8 is an uncrosslinked control. Cross-linked cell lysates were immunoblotted using CtBP2 antibody. Results of a representative experiment from three independent experiments are shown. **C**. Densitometric analysis of **B** was performed using ImageJ software and % CtBP2 dimers were quantified. **D**. OVCAR3 cells were pretreated with GMX1778 1nM for 24 h, followed by addition of increasing concentrations of JW-98 for 6 days. Cell viability was assessed *via* crystal violet staining and absorbance of solubilized dye was recorded at 590 nm. N=3; all pairwise comparisons were made using student’s t-test. * indicates p < 0.05. Error bars indicate +/- 1.0 SD.

### NAD depletion sensitizes HGSOC cells to CtBP inhibitor induced cell death

To determine if disruption of CtBP oligomers by the JW-98/GMX1778 combination translates into cellular cytotoxic effects, we pre-treated OVCAR3 HGSOC cells with GMX1778 (sub-optimal concentration after determining IC_50_) or vehicle for 24 h, followed by treatment with vehicle or increasing doses of JW-98 for 6 days and assessed cell viability **(Fig 4D)**. Strikingly, GMX1778, which had little effect by itself at 1 nM, strongly sensitized the cytotoxic effect of JW-98 at JW-98 concentrations well below the single agent IC_50_ (**Fig. 4D**). Indeed, in the presence of a non-toxic dose of GMX1778, there was a ∼3-fold loss of viability (75% to 25%) (**Fig. 4D**).

Likewise, pre-treatment of OVCAR8 and OVCA429 cells with GMX1778 strongly accentuated JW-98 cytotoxicity (**Fig. S2**).

### NAMPT inhibitor GMX1778 potentiates anti-cancer effect of JW-98 both in vitro and in vivo

To establish the in vivo efficacy of the CtBPi/NAMPTi combination, we subcutaneously xenografted immunodeficient NSG mice with OVCAR3 cells and treated the mice 3X/week for 2 weeks by oral gavage with vehicle, GMX1778 (30 mg/kg), JW-98 (100 mg/kg), or the combination of JW-98 and GMX1778, and tumor volume was measured on Days 0, 7, and 14 (**Fig. 5A**). The mice suffered no visible ill effects over the 14-day course of treatment and their weights remained stable (**data not shown**). GMX1778 treatment alone only modestly reduced tumor volume ∼35%, as did JW 98 alone, though these modest effects did not achieve statistical significance (**Fig. 5B**). However, combination therapy with GMX1778/JW-98 caused a significant 70% reduction in mean tumor volume by Day 14 (p<0.01; **Fig. 5B**), consistent with the highly cooperative effects of this drug combination observed in cultured HGSOC cells **(Fig. 4)**.

**Fig. 5.**
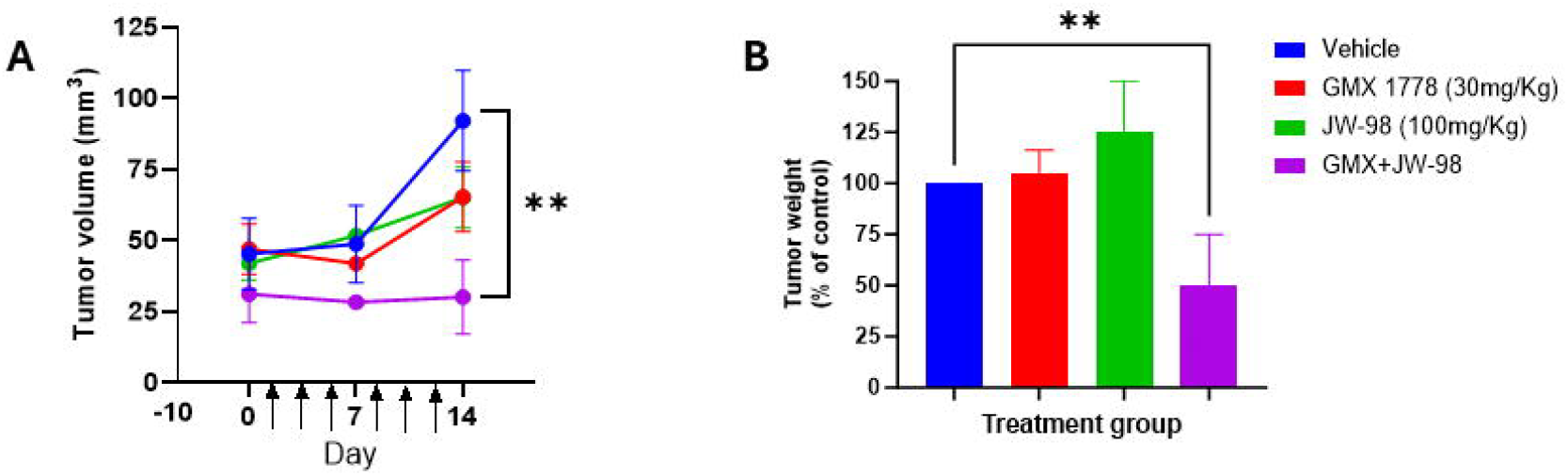
Combined CtBP/NAD inhibition abrogates growth of HGSOC xenografts. **A**. 1×10^6^ OVCAR3 cells were injected subcutaneously in the flank of NSG mice and tumors were allowed to grow for 10 days, after which animals were treated by oral gavage 3x/week for 2 weeks (indicated by arrows) with Vehicle (corn oil), GMX (30 mg/kg), JW-98 (100 mg/kg) or combination of GMX1778 and JW-98. A. Tumor volumes measured weekly. **B**. Tumor weight normalized to Vehicle-treated group and % change in tumor weight was calculated. N=4/group; Paired student t-test performed. **p < 0.01. Error bars indicate +/- 1.0 SD.

## Discussion

The efficacy of HGSOC treatment is often limited by chemoresistance. In addition, HGSOC is a complex disease driven by a combination of tumor suppressor loss and numerous non mutated oncogenic dependencies, such as overexpression of CtBP (4). We show, for the first time, overexpression of CtBP proteins in primary and recurrent/metastatic HGSOC tumors, especially CtBP2, which increases expression late in the disease. Based on the ubiquitous expression of CtBP1/2 in advanced/refractory disease, we tested a potent 3^rd^ generation CtBP dehydrogenase inhibitor, JW-98, in a panel of genetically validated HGSOC cells.

The CtBP transcriptional co-regulators are reported to dimerize or oligomerize in the presence of NAD/NADH to form transcriptional complexes regulating their oncogenic activities (16). NAD is a key metabolite required for normal cellular growth and proliferation; however, tumor cells are in greater need of NAD than normal cells to leverage the ever-growing demand for cellular ATP needed for proliferation and growth (17). We strategized that limiting NAD supply to tumor cells using the NAMPT inhibitor GMX1778 would further abrogate CtBP’s oncogenic activity and lead to cell death. Indeed, our earlier work in pancreatic cancer has shown that limiting NAD synthesis coupled with CtBP inhibition improves the cytotoxic efficacy of CtBPi by disruption of the dimeric CtBP complex (8). Indeed, when we limited NAD synthesis in HGSOC cells using a NAMPT inhibitor (GMX1778), the combination of GMX1778 and JW-98 disrupted CtBP dimerization, a key step in its oncogenic activity. NAD depletion was synergistically cytotoxic with JW-98, leading to enhanced cell death in vitro. Combined treatment of mice with GMX1778 and JW-98 safely abrogated HGSOC xenograft tumor growth while neither drug alone was effective.

HGSOC is a lethal cancer in women due to the lack of effective systemic therapies in advanced, especially chemo-resistant, disease. Overall, our findings strongly support the combined inhibition of NAD/CtBP as an innovative strategy to target HGSOC. Furthermore, our *in vivo* results of the OVCAR3 xenograft study using the combination of JW-98 and GMX1778 show promising therapeutic utility that should be explored further in additional models, including patient-derived xenografts, to determine the broad applicability of this therapeutic strategy in HGSOC.

