## Supplementary figures and images for "Combined inhibition of NAD synthesis and C-terminal binding protein cooperatively induce cell death and inhibit growth of High Grade Serous Ovarian Carcinoma"

### Figure S1

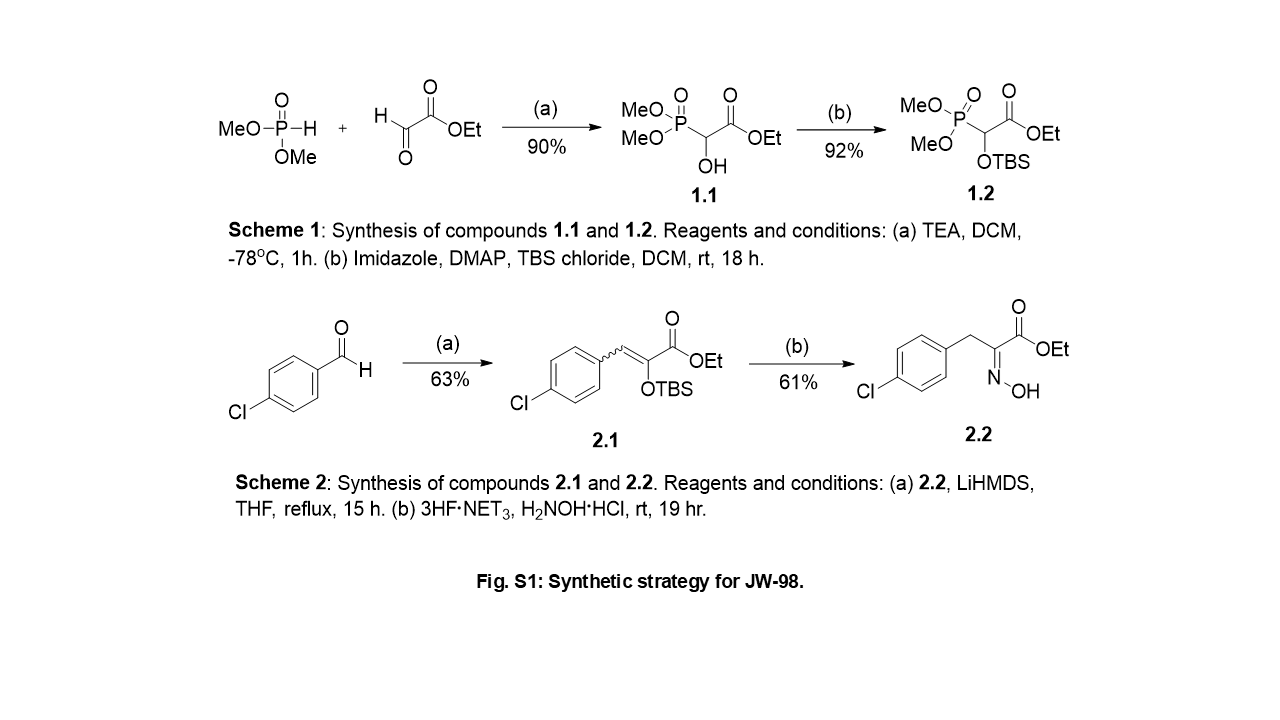

### Figure S2

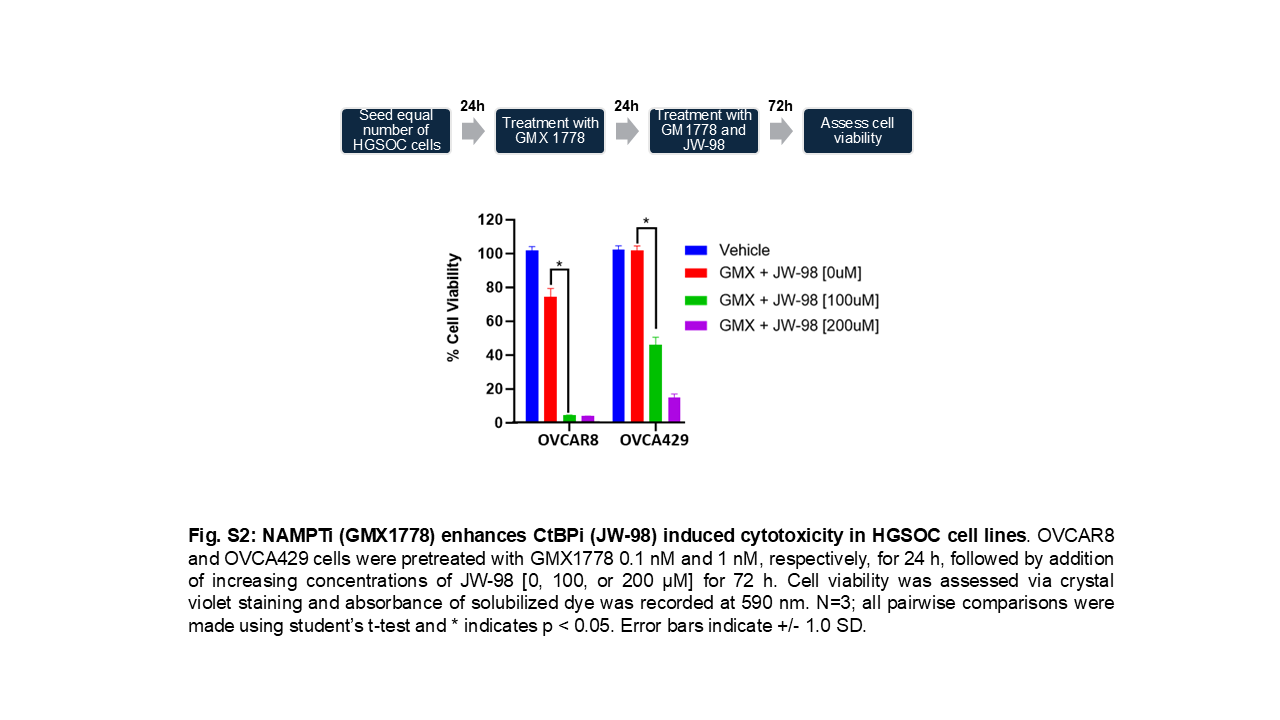
